# Comprehensive biodiversity analysis via ultra-deep patterned flow cell technology: a case study of eDNA metabarcoding seawater

**DOI:** 10.1101/515890

**Authors:** GAC Singer, N Fahner, J Barnes, A McCarthy, M Hajibabaei

## Abstract

The characterization of biodiversity is a crucial element of ecological investigations as well as environmental assessment and monitoring activities. Increasingly, amplicon-based environmental DNA metabarcoding (alternatively, marker gene metagenomics) is used for such studies given its ability to provide biodiversity data from various groups of organisms simply from analysis of bulk environmental samples such as water, soil or sediments. The Illumina MiSeq is currently the most popular tool for carrying out this work, but we set out to determine whether typical studies were reading enough DNA to detect rare organisms (i.e., those that may be of greatest interest such as endangered or invasive species) present in the environment. We collected sea water samples along two transects in Conception Bay, Newfoundland and analyzed them on the MiSeq with a sequencing depth of 100,000 reads per sample (exceeding the 60,000 per sample that is typical of similar studies). We then analyzed these same samples on Illumina’s newest high-capacity platform, the NovaSeq, at a depth of 7 million reads per sample. Not surprisingly, the NovaSeq detected many more taxa than the MiSeq thanks to its much greater sequencing depth. However, contrary to our expectations this pattern was true even in depth-for-depth comparisons. In other words, the NovaSeq can detect more DNA sequence diversity within samples than the MiSeq, even at the exact same sequencing depth. Even when samples were reanalyzed on the MiSeq with a sequencing depth of 1 million reads each, the MiSeq’s ability to detect new sequences plateaued while the NovaSeq continued to detect new sequence variants. These results have important biological implications. The NovaSeq found 40% more metazoan families in this environment than the MiSeq, including some of interest such as marine mammals and bony fish so the real-world implications of these findings are significant. These results are most likely associated to the advances incorporated in NovaSeq especially patterned flow cell, which prevents similar sequences that are neighbours on the flow cell (common in metabarcoding studies) from being erroneously merged into single spots by the sequencing instrument. This study sets the stage for incorporating eDNA metabarcoding in comprehensive analysis of oceanic samples in a wide range of ecological and environmental investigations.

## 2 Introduction

The inventorying and monitoring of biological diversity is a fundamental component of ecological and environmental studies. Additionally, characterizing biodiversity is part of the environmental impact assessments and ongoing environmental monitoring that are required by industry operating in environmentally sensitive locations ^1^. Stakeholders are increasingly becoming more concerned about environmental stewardship, and this applies equally to the terrestrial ^2^, freshwater ^3^, and marine environments ^4^ and covers all major taxonomic groups. A recent United Nations conference (UN Biodiversity Conference, Egypt, November 2018), highlighted the need for monitoring and protecting biodiversity with the key message of “investing in biodiversity for people and planet”. Despite the extreme importance of these efforts, the technology for carrying out biodiversity assessments has remained static for decades, relying heavily on observational data and capturing whole organisms from their environment for morphological analysis. Unfortunately, these procedures are error-prone, time-consuming, expensive, and tend to ignore small but ecologically important flora and fauna simply because they’re difficult to identify visually ^2,5^.

Over the past decade, increasing attention has been paid to the analysis of environmental DNA (eDNA)—a combination of DNA from whole cellular material or that is shed from organisms as they move through their environment. The existence of large reference databases (especially the common “DNA barcode” marker, cytochrome oxidase c subunit 1, or COI^6,7^) with the power of modern DNA sequencing instruments, enables environmental metabarcoding—the identification of many individual species from a simple water or sediment sample. Environmental metabarcoding is much faster than conventional techniques, is less labour-intensive, does not rely on the expertise of taxonomists, and produces orders of magnitude more information ^8^.

Many side-by-side comparisons have been made between traditional morphological assessments and eDNA-based assessments. In all cases, eDNA is capable of detecting far more taxa overall. However, many of these studies also find that some organisms detected by traditional methods in the environment fail to be detected through metabarcoding ^1,4,9–13^ There are a number of potential reasons for this discrepancy: the use of “universal” primers that don’t amplify some taxa as well as others ^5^; employing markers that have biased representation in reference databases ^14^; or an inadequate depth of sequencing to detect eDNA that is in low abundance. These factors are especially important when eDNA analyses are performed to track specific target organisms that might be present in low abundance in complex settings such as the oceans (e.g., endangered species or invasive species).

The goal of the present study was to investigate the influence of sequencing depth and more advanced workflow, including a patterned flowcell, offered by illumina’s NovaSeq platform on the ability to detect biological diversity present in a sample. To carry out this work, we analyzed samples on an Illumina MiSeq instrument at a sequencing depth that is typical of similar studies (Table 1), then analyzed those same PCR products on an Illumina NovaSeq 6000—the most advanced HTS instrument available today— with a sequencing depth approximately 700 times greater than that of the MiSeq. Not surprisingly, the NovaSeq detected many more taxa than the MiSeq: specifically, with one marker the NovaSeq detected 200% more metazoan families than the MiSeq. Contrary to our expectations, the NovaSeq still outperformed the MiSeq even when we subsampled the data to make depth-for-depth comparisons, suggesting that the NovaSeq has superior qualities beyond its much greater sequencing capacity.

**Table 1:** A survey of recently published metabarcoding studies shows that the Illumina MiSeq is the most commonly used instrument to analyze these data, and that sequencing depth per sample varies widely but has a median of approximately 60,000 reads. Cases where the sequencing depth was not directly reported and had to be inferred indirectly are indicated with an asterisk.

| Reference | Publication type | Substrate | Type of study | Sequencing platform | Average raw reads per |
| --- | --- | --- | --- | --- | --- |

|  |  |  |  |  | **amplicon**<br>**per sample** |
| --- | --- | --- | --- | --- | --- |
| 29 | Pre-print | Leech gut contents | Targeted (vertebrates) | Illumina MiSeq | Unknown |
| 30 | Peer-reviewed | Herbivore feces | General biodiversity | IonTorrent PGM | 4,156 (post-filtering) |
| 31 | Peer-reviewed | Soil | Targeted (*Phytophthora*) | Roche GS Junior | Unknown |
| 32 | Peer-reviewed | Freshwater ecosystem | Targeted: planktonic protists | Illumina MiSeq | 150,000* |
| 33 | Peer-reviewed | Turbid freshwater | Targeted (vertebrates) | Illumina MiSeq | 7,181 (post-filtering) |
| 34 | Peer-reviewed | Sea water | General biodiversity | Roche GS-FLX | 21,192* (post-filtering) |
| 35 | Pre-print | Brackish water | General biodiversity | Illumina MiSeq | 200,185 (post-filtering) |
| 36 | Peer-reviewed | High-salinity lake | General biodiversity | Illumina HiSeq 2500 | 124,779 for bacteria<br>~37,000 for archaea and eukaryota (post-filtering) |
| 37 | Pre-print | Wheat and oilseed rape residues | General biodiversity | Illumina MiSeq | 80,000* (raw) |
| 38 | Peer-reviewed | Scat | Targeted (truffle fungi) | Illumina MiSeq | <60,000* |
| 39 | Peer-reviewed | Marine sediment | Targeted (microbial) | Illumina MiSeq | 35,663 |
| 40 | Pre-print | Mock community | Targeted (Symbiodiniaceae) | Illumina MiSeq | 6,309* (raw) |
| 41 | Peer-reviewed | Plant leaves | Targeted (fungi) | Illumina MiSeq | 112,931* |
| 42 | Peer-reviewed | Freshwater | General diversity | IonTorrent PGM | 3,978 (post-filtering) |
| 43 | Peer-reviewed | Shrimp stomach contents | General biodiversity | Illumina MiSeq | 350,000* |
| 44 | Peer-reviewed | Soil | General biodiversity | Illumina MiSeq | 61,256 (raw) |
| 45 | Peer-reviewed | Bat guano | General biodiversity | Illumina MiSeq | 106,858 |
| 46 | Peer-reviewed | Bee nest chambers | General biodiversity | Illumina MiSeq | 4,736 (post-filtering) |
| 47 | Peer-reviewed | Sea water | Targeted (dinoflagellates) | Roche GS Junior | 252,086* (raw) |
| 48 | Peer-reviewed | Soil | General biodiversity | Illumina MiSeq and HiSeq | 880,000* (raw) |

## 3 Methods

### 3.1 Sample collection

Triplicate 250mL water samples were taken from surface water simultaneously. Samples were taken from eight locations along two transects in Conception Bay, Newfoundland and Labrador, Canada, on October 13-14, 2017 (Figure 1). These samples cover a range from near-shore to approximately 10km offshore (with a sea bottom depth ranging from a few metres nearshore to approximately 200 metres in the middle of the bay).

**Figure 1:**
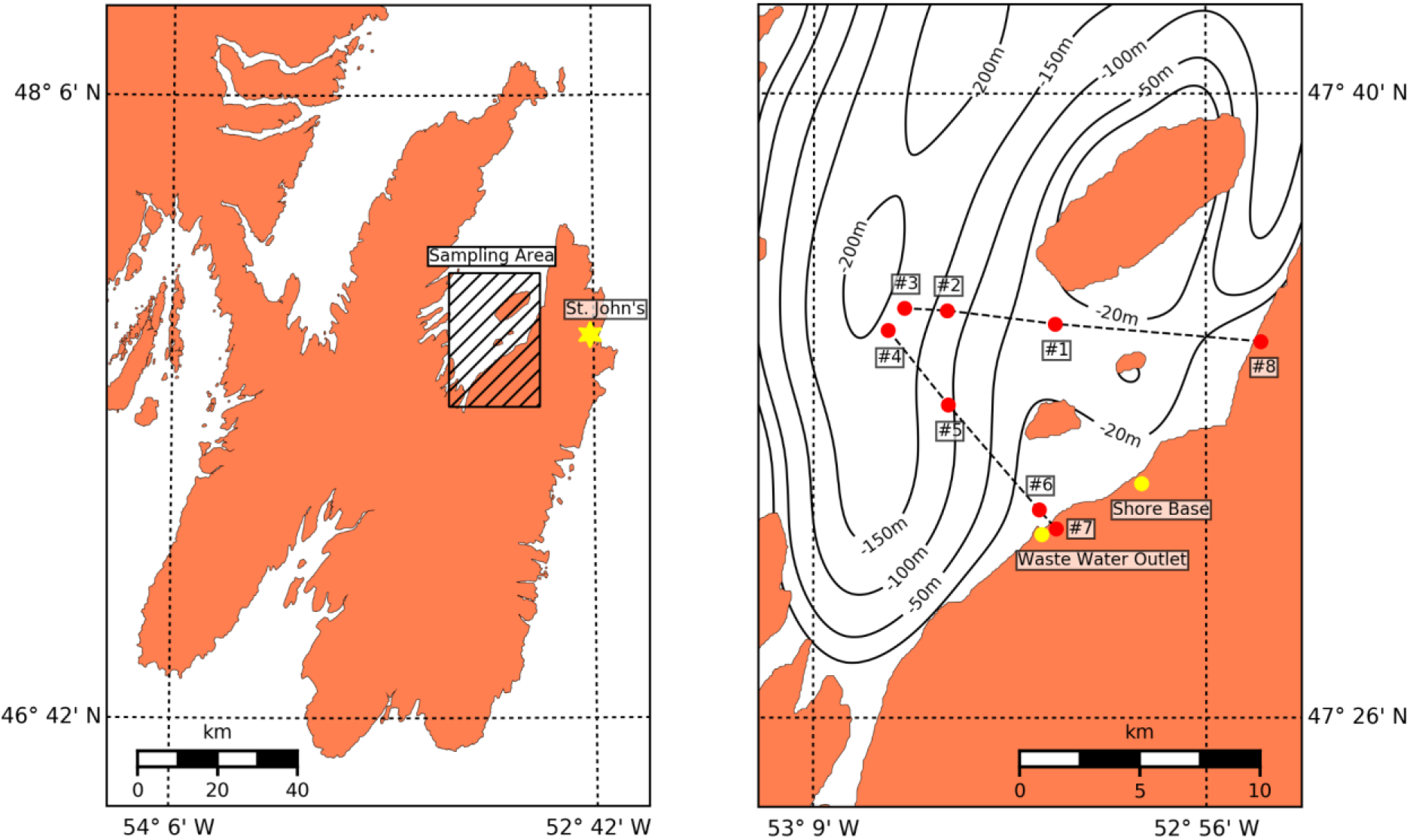
Location of the eight sampling sites from Conception Bay, Newfoundland and Labrador, Canada.

### 3.2 Laboratory procedures

#### 3.2.1 DNA extraction

Filtration and DNA extraction was done in PCR clean laminar flow hoods (AirClean Systems) thoroughly decontamination with ELIMINase (Decon Labs) and 70% EtOH prior to each set of three sample replicates. Water samples were thawed at 4°C and immediately filtered with 0.22 μm PVDF Sterivex filters (MilliporeSigma). The DNeasy PowerWater Kit (Qiagen) was used to extract DNA with the automated QIAcube platform (Qiagen), following the DNeasy PowerWater IRT protocol. For lysis, bead tubes were heated for five minutes at 65°C and then vortexed for ten minutes. Negative controls were generated during filtration and extraction to screen for contamination and cross-contamination. Filtration and extraction were done in a pre-PCR room isolated from post-PCR rooms.

#### 3.2.2 DNA library preparation

Two fragments were amplified by PCR from the 5’ end of the standard COI barcode region: the 235 bp F230 fragment^18^, and the 232 bp Mini_SH-E fragment^15^. Illumina-tailed PCR primers (tails underlined) were used to amplify targets: The F230 forward primer (LC01490; 5’<u>-TCG TCG GCA GCG TCA GAT GTG TAT AAG AGA CAG</u> GGT CAA CAA ATC ATA AAG ATA TTG G-3’; ^16^), the Mini_SH-E forward primer (Fish_miniE_F_t; 5’<u>-TCG TCG GCA GCG TCA GAT GTG TAT AAG AGA CAG</u> ACY AAI CAY AAA GAY ATI GGC AC-3’), and the reverse primer (230_R/Fish_miniE_R_t; 5’<u>-GTC TCG TGG GCT CGG AGA TGT GTA TAA GAG ACA</u> GCT TAT RTT RTT TAT ICG IGG RAA IGC-3’) was used for both F230 and Mini_SH-E fragments. Each amplification reaction contained 0.6 μL DNA, 1.5 μL 10X reaction buffer, 0.6 μL MgCl2 (50 mM), 0.3 μL dNTPs mix (10 mM), 0.3 μL of each Illumina-tailed primer (10 μM), and 0.3 μL Platinum Taq (0.5 U/μL; Invitrogen) in a total volume of 15 μL. PCR conditions were initiated with a heated lid at 95°C for 3 mins, followed by 35 cycles of 94°C for 30 s, 46°C for 40 s, and 72°C for 1 min, and a final extension at 72°C for 10 mins. Three PCR replicates were amplified from each sample with the ProFlex thermocycler (Thermo Fisher) and then pooled for a single PCR cleanup with the QIAquick 96 PCR purification kit (Qiagen; 60 μL elution volume). Agarose (1.5% w/v) gel electrophoresis was used to verify amplification of samples, and for quality control of negative controls from PCR, extraction, filtration, and field collection. Each indexing reaction contained 2 μL amplicon DNA, 2.5 μL 10X reaction buffer, 1 μL MgCl2 (50 mM), 0.5 μL dNTPs mix (10 mM), 1 μL of F indexing primer (5 μM), 1 μL R indexing primer (5 μM), and 0.5 μL Platinum Taq (0.5U/μL; Invitrogen) in a total volume of 25 μL. Unique dual Nextera indexes were used to mitigate index misassignment (IDT; 8-bp index codes). PCR conditions were initiated with a heated lid at 95°C for 3 mins, followed by 12 cycles of 95°C for 30 s, 55°C for 30 s, and 72°C for 30 s, and a final extension at 72°C for 5 mins. Indexing success was verified on the Bioanalyzer (Agilent) with the DNA 7500 kit. Samples were quantified with Quant-iT PicoGreen dsDNA assay with a Synergy HTX plate fluorometer (BioTek) and pooled to normalize DNA concentration. Libraries were cleaned with three successive AMPure XP cleanups: Left side selection with bead:DNA ratios of 1X, then 0.9X, and a right-side selection with 0.5X. Libraries were quantified with a Qubit fluorometer (Thermo Fisher) and the size distribution was checked with the DNA 7500 kit on the Agilent 2100 Bioanalyzer. Two libraries containing F230 or FishE amplicons from nine samples were sequenced on the Illumina MiSeq with two 600-cycle v3 kits. Two libraries containing F230 or FishE amplicons from 24 samples were pooled with other libraries and sequenced with two MiSeq 600-cycle v3 kits. Field, filtration and extraction negatives were also sequenced in these MiSeq runs. Two libraries containing F230 and FishE amplicons were sequenced with a 300-cycle S4 kit on the NovaSeq 6000 following the NovaSeq XP workflow.

#### 3.2.3 Bioinformatics

We employed two different workflows to analyze our data in order to reduce the possibility that our results were an artefact of the method used. In both workflows, base calling and demultiplexing were performed using Illumina’s bcl2fastq software (version 2.20.0.422). Primers were then trimmed from the forward and reverse reads using cutadapt v1.16 ^17^ with the default error tolerance and a minimum overlap equal to half the primer length. We discarded read pairs in which the primer was missing from either the forward or reverse read. After this stage the two methodologies diverged and are described separately below.

##### 3.2.3.1 DADA2 workflow

DADA2 v1.8.0 ^18^ was used to perform quality filtering and joining of paired reads (maxEE=2, minQ=2, truncQ=2, maxN=0), and denoising (using default parameters) to produce exact sequence variants (ESVs). This was performed independently on the MiSeq and NovaSeq data since their error patters are presumed to be different and therefore they require different models to be trained. Singleton ESVs were discarded. To rapidly evaluate the overlap in ESVs between the two instruments, MD5 hashes ^19^ were generated for each of the ESV sequences and then these sets of hashes were compared between the MiSeq and the NovaSeq.

##### 3.2.3.2 OTU clustering workflow

Vsearch v2.8.2 ^20^ was first used to join the paired ends of the reads (using default parameters), and perform quality filtering (using default parameters). The reads for the NovaSeq and MiSeq were then dereplicated, and these reads were combined into a single file so that OTU clustering (using the cluster_fast setting) could be performed on the entire set using an identity threshold of 97%. As with the ESVs, singleton OTUs were discarded.

##### 3.2.3.3 Taxonomic assignment

NCBI’s blastn tool v2.6.0 ^21^ was used to compare ESV sequences against the nt database (downloaded August 2018), using an e-value cut-off of 0.001. We filtered the resulting hits with the requirement of having at least 90% identities across at least 90% of the query sequence. In cases where there was not a single unambiguous best hit, we used a majority consensus threshold of 80% to assign taxonomy ^22^.

##### 3.2.3.4 Accumulation curves

Original read memberships were tracked through the various analytical steps: dereplication followed by OTU clustering, or ESV generation using DADA2. Subsamples were then generated using sampling proportional to the original read abundances with the “choices” function within the Python programming language’s “random” module ^23^. These reads were then mapped to their respective ESVs/OTUs for comparison between the two DNA sequencing platforms.

##### 3.2.3.5 Availability of data

All data have been deposited into NCBI’s Sequence Read Archive under accession number PRJNA513845.

## 4 Results

### 4.1 The Illumina MiSeq is currently the most popular metabarcoding platform

Twenty of the most-recently indexed papers in Google Scholar featuring the “metabarcoding” keyword were obtained in early November 2018 to perform a mini-metanalysis of the instrument most such studies are favouriting at the moment, as well as the depth of sequencing per sample that is generally employed. As shown in Table 1, the Illumina MiSeq is by far the instrument of choice presently, having been used by 14 (70%) of these studies. Sampling depth was not always reported clearly but was inferred where possible. Among these studies there was an extremely wide variance in sequencing depth, ranging from less than 10,000 reads per sample to nearly 900,000. However, the median was 60,000 (with a median absolute deviation of 55,000).

### 4.2 The NovaSeq finds more ESVs per sample than the MiSeq, even at the same sequencing depth

Based on our literature survey (Table 1), we decided to analyze our own samples on the MiSeq with a targeted sequencing depth of 100,000 reads per amplicon per sample—approximately 50% greater than the median sequencing depth of those studies. Two amplicons were analyzed, FishE and F230, both located with the standard barcode region of the mitochondrial gene cytochrome oxidase C subunit 1 (COI). Post-filtering, the mean depth per sample was 118,290 reads for the FishE marker and 84,500 for the F230 marker. We then analyzed these same PCR products on the Illumina NovaSeq 6000 at much greater depth, averaging 7 million reads per amplicon per sample. The resulting reads were processed using the DADA2 pipeline as described in the Methods. Perhaps not surprisingly given the ~700x greater sequencing depth, the NovaSeq was able to find more exact sequence variants (ESVs) in each sample than the MiSeq. To our surprise, however, even after rarefying the NovaSeq data to match the sequencing depth of the MiSeq, we still found greater diversity (i.e., more ESVs) within the NovaSeq data for the FishE (Figure 2) and F230 (Supplementary Figure 1) amplicons. Moreover, while there was substantial overlap between the ESVs found between the two platforms, the MiSeq had very few ESVs unique to itself while the NovaSeq found many ESVs that the MiSeq missed. We highlight that the exact same PCR products were used for both instruments, so these results cannot be the consequence of stochastic PCR biases. The two sites with higher diversity—7 and 8—are located close to shore and site 7 in particular is close to a wastewater outlet (Figure 1).

**Figure 2:**
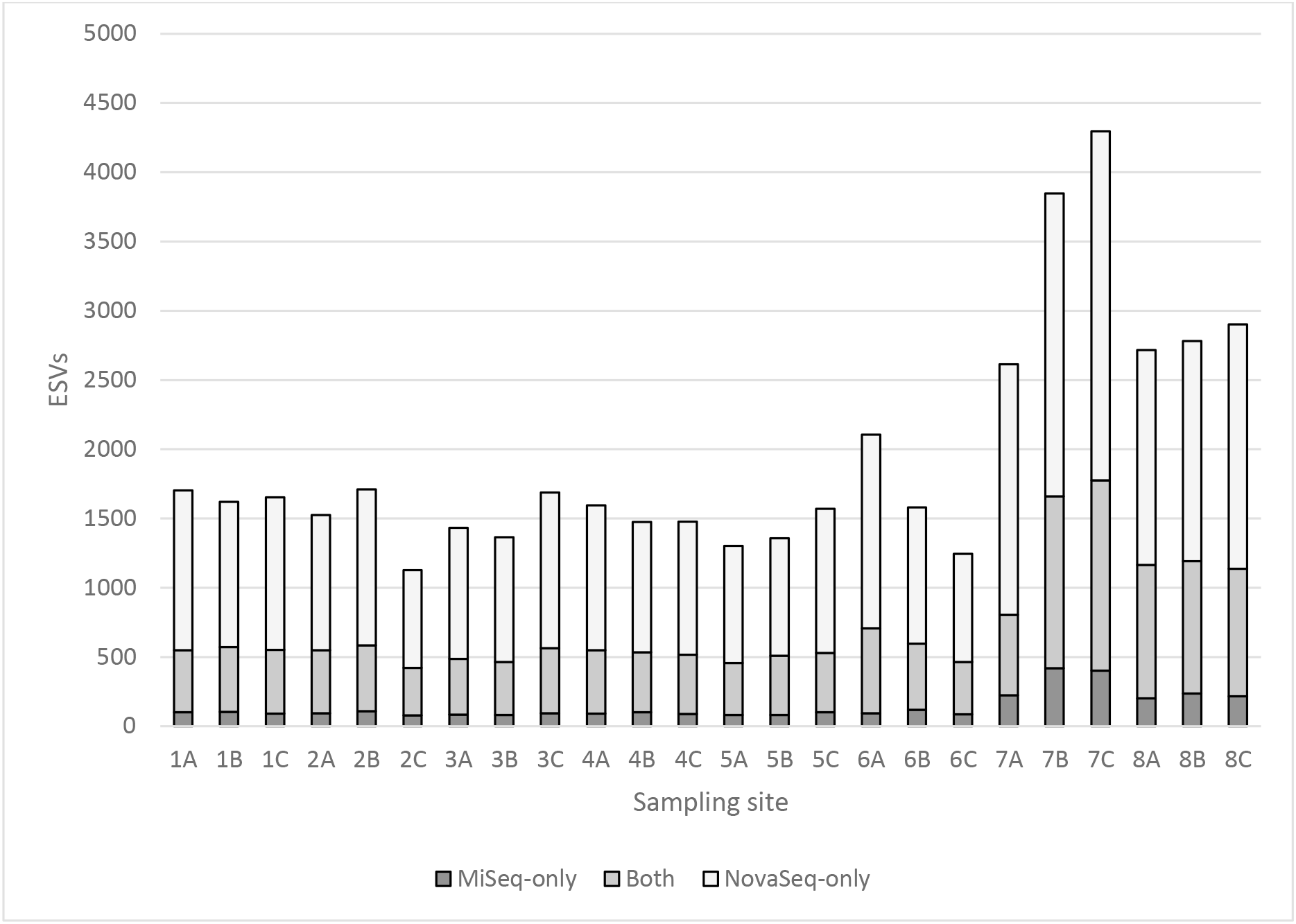
For each biological replicate (A, B, C) within each sampling site (1 to 8), the NovaSeq (light bars) was able to find a greater number of ESVs than the MiSeq (dark bars) even when the NovaSeq data are subsampled to match the sequencing depth of the MiSeq. Sites 7 and 8 are very close to shore, and site 7 in particular is near a wastewater outlet.

### 4.3 This trend is even more pronounced when plotted as an accumulation curve

When we combined all samples and then performed subsampling experiments to generate accumulation curves, the ability of the NovaSeq to detect new ESVs becomes even more stark: at each simulated sequencing depth, the NovaSeq detects greater biological diversity (i.e., ESVs) than the MiSeq (Figure 3 for the FishE amplicon; see Supplementary Figure 2 for the F230 amplicon). Curiously, while greater depth seems to reveal increasing numbers of ESVs on the NovaSeq (even beyond 2.5 million reads/sample), it is not clear that greater depth adds any new information for the MiSeq: the number of ESVs detected appears to level off at approximately 5,000. This strongly suggests that the NovaSeq outperforms the MiSeq in a depth-independent manner.

**Figure 3:**
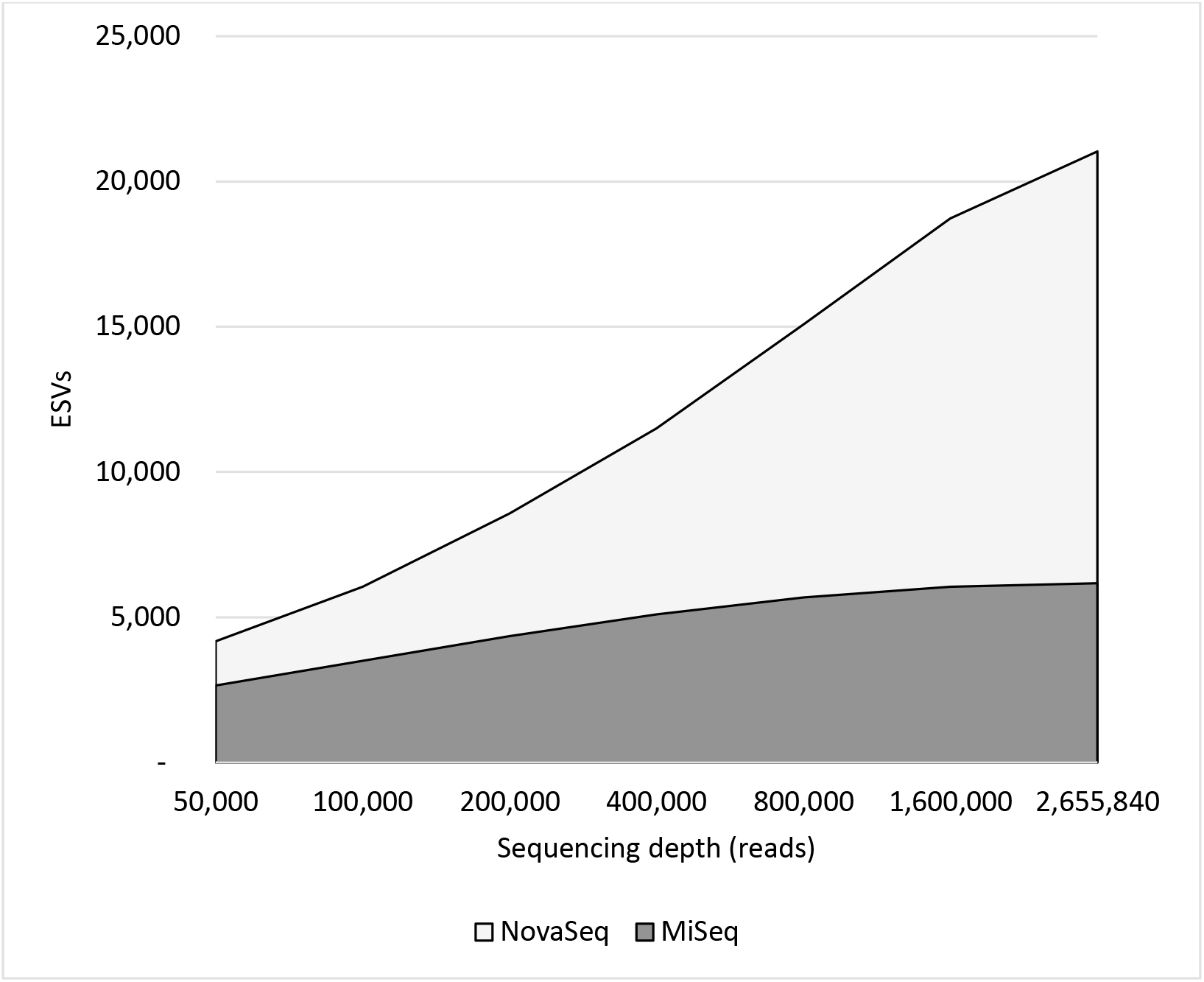
Accumulation curves generated by subsampling the MiSeq (dark curve) and NovaSeq data (light curve) for the FIshE amplicon shows that depth-for-depth the NovaSeq detects greater biological diversity than the MiSeq

### 4.4 This trend is not an artefact of the DADA2 error-correcting model

DADA2 generates ESVs by applying an error correction model to raw FASTQ files, attempting to fix errors that were introduced through PCR or sequencing ^18^. However, while the MiSeq reports base call qualities using pseudo-continuous Phred scores that can range from 0-40, the NovaSeq’s FASTQ files bin qualities into just four levels^24^. We therefore suspected that the phenomenon we were observing might be an artefact of the DADA2 program. Specifically, we hypothesized that the algorithm might be under-correcting errors in the NovaSeq data leading to a spurious increase in the number of ESVs. For this reason we repeated our analysis with simple OTU clustering at a 97% similarity threshold (described in greater detail in the Methods). OTU clustering applies no error correction model at all and is simply based on sequence similarity measures, and should therefore have the same performance on NovaSeq data as it does on MiSeq data. To our surprise, when accumulation curves were generated to compare the two instruments depth-for-depth, the NovaSeq once again outperformed the MiSeq (Figure 4).

**Figure 4:**
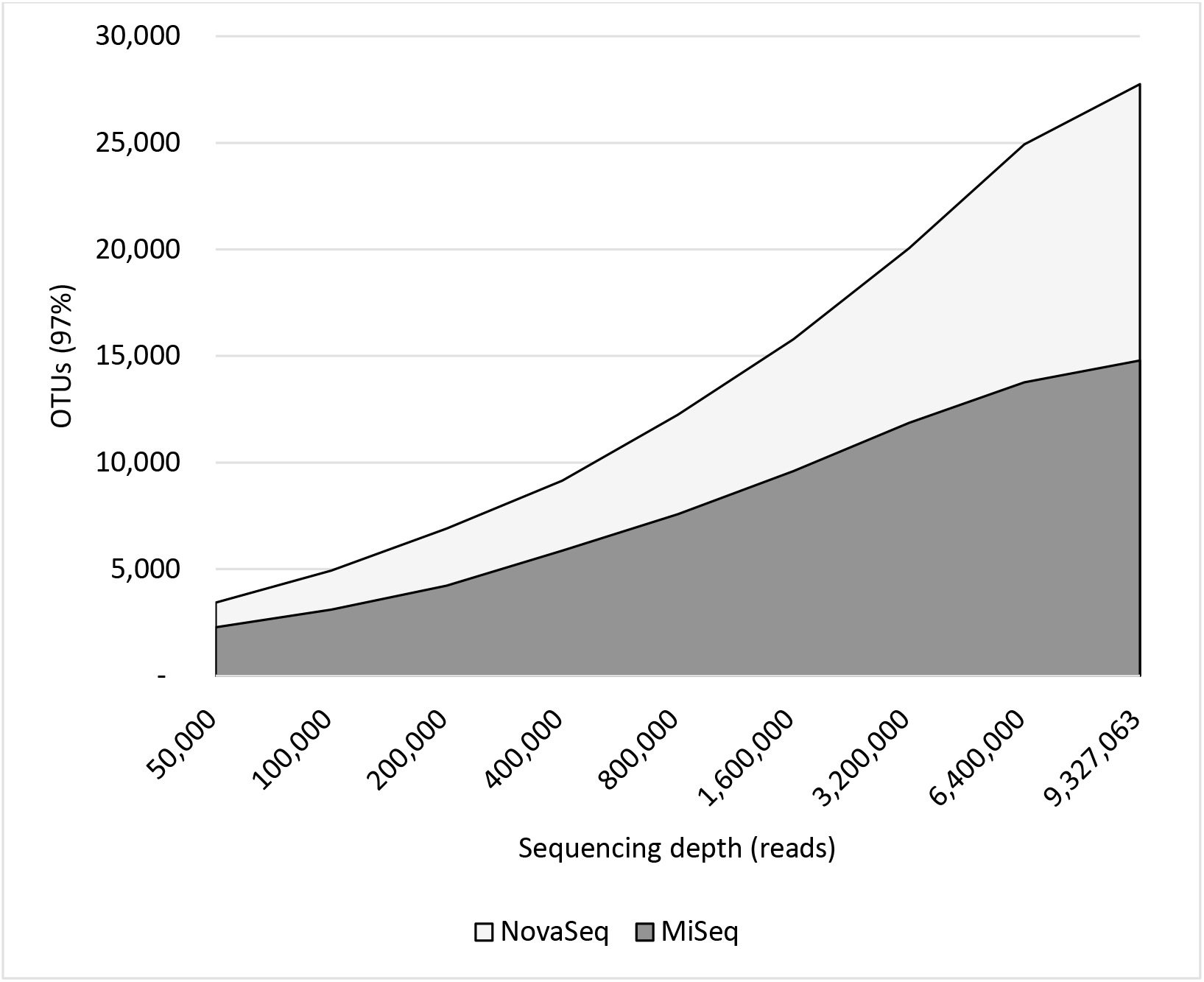
Accumulation curves of MiSeq (dark curve) and NovaSeq data (light curve) based on OTU clustering with a 97% identity threshold. At each sequencing depth, the NovaSeq finds more OTUs than the MiSeq, similar to the ESV data.

We note that Figure 3 and Figure 4 look quite different from each other in two ways: (1) the number of OTUs detected is far greater than the number of ESVs; and (2) while the number of new ESVs levels out for the MiSeq in Figure 3, the trajectory continues upward for the OTUs in Figure 4. This is due to the very different methodologies employed to generate ESVs versus OTUs. Since OTU clustering makes no attempt to model and correct for PCR and sequencing errors, the raw number of OTUs is expected to be much greater than the number of ESVs detected—many OTUs are simply the product of the accumulation of errors. By similar reasoning, both the MiSeq and NovaSeq OTU curves continue to rise with greater sequencing depth because additional sequencing errors will be encountered with that greater depth.

### 4.5 Greater sequencing depth on the MiSeq cannot achieve the level of diversity detected on the NovaSeq

The MiSeq’s accumulation curve in Figure 3 suggests that additional sequencing depth would not increase the number of ESVs detected, but to thoroughly test this point we re-ran three samples (sites 1, 3, and 6, each with three biological replicates for a total of 9 samples) on the MiSeq at much greater sequencing depth—approximately 1 million reads per amplicon per sample—and then compared these data to the NovaSeq data. As illustrated for the FishE amplicon in Figure 5, adding this greater sequencing depth to the MiSeq only marginally improves its detection of diversity from the samples. Conversely, the NovaSeq continues to detect new ESVs with greater sequencing depth. Note that the total number of ESVs detected is lower than that of Figure 3, but this is because of the smaller number of samples analyzed in this experiment (three sites versus eight). Again, we repeated this experiment with the F230 amplicon and found the same trend (see Supplementary Figure 3).

**Figure 5:**
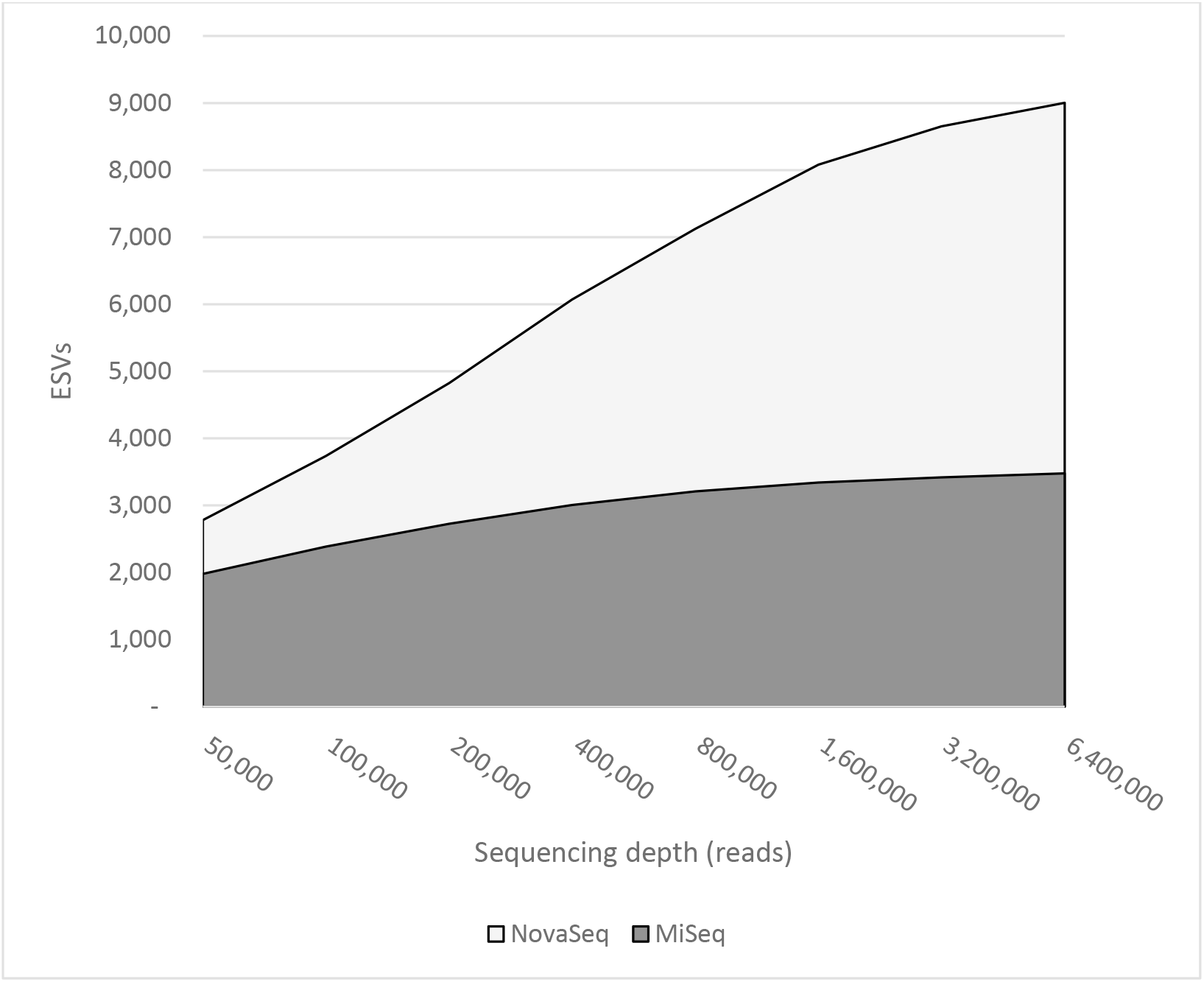
Re-running a subset of our samples on the MiSeq (dark curve) with much greater sequencing depth only added a very small number of new ESVs. Conversely, greater sequencing depth continues to add new ESVs to the NovaSeq data (light curve).

Figure 5 shows that even beyond 5 million reads the NovaSeq was still finding new ESVs with no sign of plateauing. As before, we suspected this might be the result of the DADA2 algorithm under-correcting sequencing errors in the NovaSeq data. To examine this possibility, we ran the accumulation curve out to its maximum and found that the curve does indeed hit a plateau of just over 9,000 ESVs at a sequencing depth of approximately 10 million reads (Figure 6). This result indicates that the pattern observed for NovaSeq data are not likely to be an artefact of the DADA2 analysis. Moreover, it indicates that extremely deep sequencing is required if one wants to have a comprehensive survey of biodiversity in a region.

**Figure 6:**
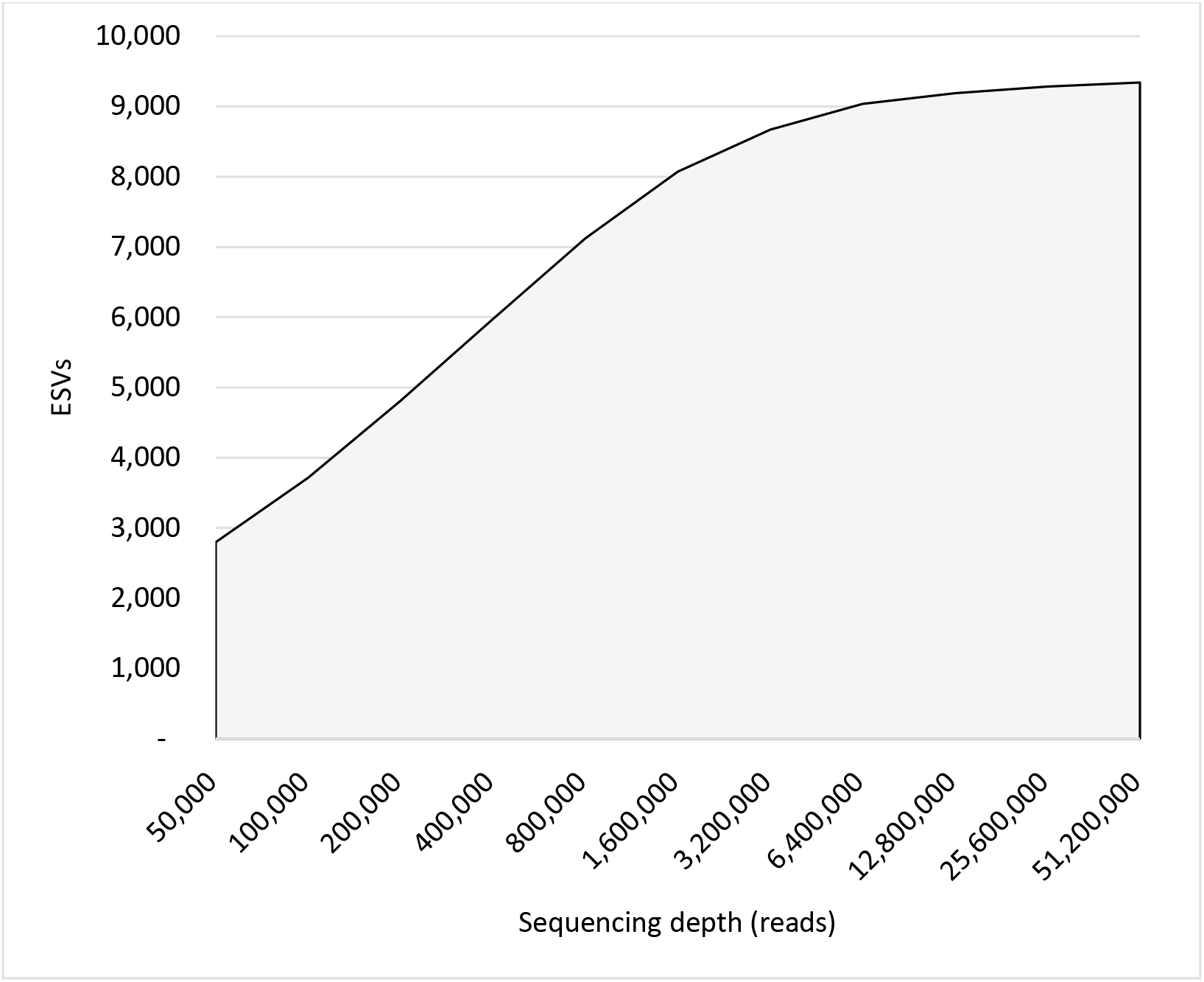
Accumulation curve of pooled NovaSeq data. The NovaSeq does eventually reach a plateau where no new ESVs are detected, albeit at an extreme sequencing depth of ~10 million reads.

### 4.6 The differences between MiSeq and NovaSeq results have biological significance

Our NovaSeq results indicate that in the locations in which we sampled, approximately 9,300 FishE ESVs are present (Figure 6). However, the MiSeq was only able to obtain approximately 3,500 ESVs even at an unrealistically-high sequencing depth (Figure 5). This suggests that the MiSeq could not identify approximately 60% of the diversity present in the environment. In order to determine taxonomic/biological breadth of these ESVs we performed taxonomic assignment on all the ESVs from both instruments—i.e., we combined the results of both MiSeq runs to give that platform the best possible chance of finding all the taxa present—and found that the MiSeq data contained 80 identifiable families. The NovaSeq also identified these same 80 families but was also able to identify an additional 32—a 40% increase. Those families unique to the NovaSeq analysis are listed in Table 2. Some of the taxa missing from the MiSeq data are of significant interest, including marine mammals (Delphinidae) and several fish. Interestingly, when these taxa are plotted on a circular dendrogram, we don’t observe any obvious phylogenetic pattern to the distribution of missing taxa on the MiSeq (Figure 7). Rather, it appears that the NovaSeq was generally able to detect more families within each order than the MiSeq.

**Table 2:**
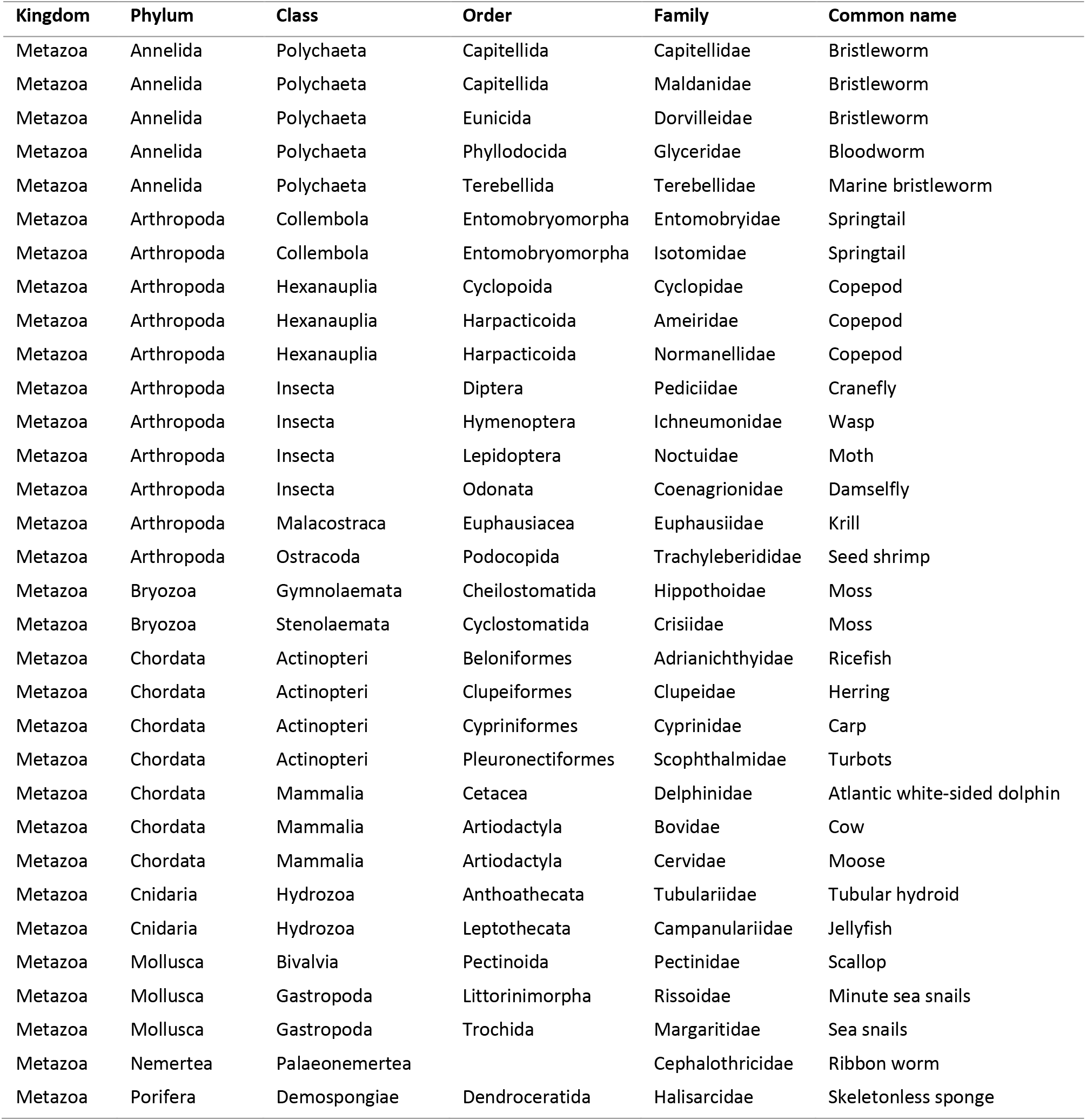
List of families detected with the NovaSeq but not with the MiSeq. Many are biologically significant taxa, including dolphins and several fish

**Figure 7:**
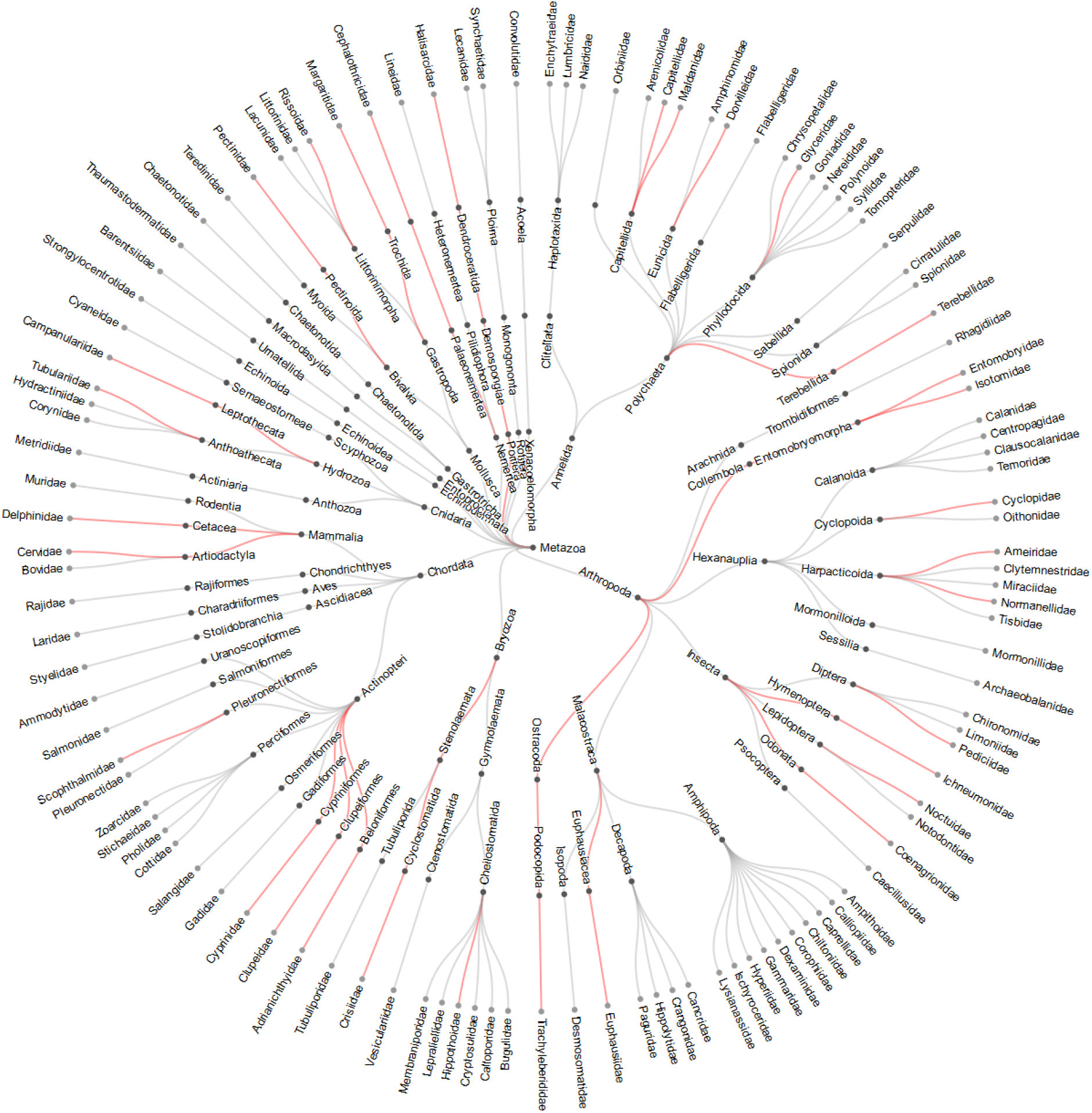
Radial dendrogram of taxa identified in our experiments. The leaves of the tree represent family-level taxa. Grey edges on the tree indicate taxa that both the NovaSeq and MiSeq platforms detected. Edges in red were only detected on the NovaSeq; there were no taxa unique to the MiSeq.

## 5 Discussion

### 5.1 5.1 Most environmental metabarcoding studies are not sequencing deep enough

Our results suggest that using seawater as the source of environmental DNA at a typical sequencing depth of 60,000 reads per sample, only half of the diversity detectable by the MiSeq will be captured (Figure 3). To reach the MiSeq’s detection limits for analysis of seawater one would have to aim for 0.8-1 million reads per sample per marker—more than ten times the typical depth of sequencing currently performed in most metabarcoding studies. We further note that the samples from our study were obtained from the North Atlantic Ocean where biodiversity is presumably far less than that which might be found in tropical regions. For this reason, even deeper sequencing may be required in regions or substrates that have very high biodiversity.

### 5.2 Even when matched for depth, the NovaSeq can detect greater diversity than the MiSeq

The most remarkable finding in this study is that the NovaSeq can detect many taxa that the MiSeq cannot—even when the depth of sequencing is matched. This is true on a PCR reaction-by-PCR reaction basis (Figure 2), and even greater depth of sequencing on the MiSeq cannot overcome this obstacle (Figure 5). The outcome is that there may be a great deal of missing biodiversity in MiSeq analyses (Figure 7).

Although we have no quantitative measurements of the abundance of taxa present in the locations we sampled, we note that many of the taxa missing from the MiSeq analysis are likely to have a very low abundance of eDNA (e.g., marine mammals, terrestrial organisms) compared to taxa where whole organisms or gametes may be present in the water samples (e.g., zooplankton). We can approximate this by looking at read abundances (Figure 8). If we assume that read abundances roughly correlate with the original biomass present^25^ then it does indeed seem that the MiSeq is less capable of sequencing this low abundance eDNA than the NovaSeq, even at very high sequencing depths.

**Figure 8:**
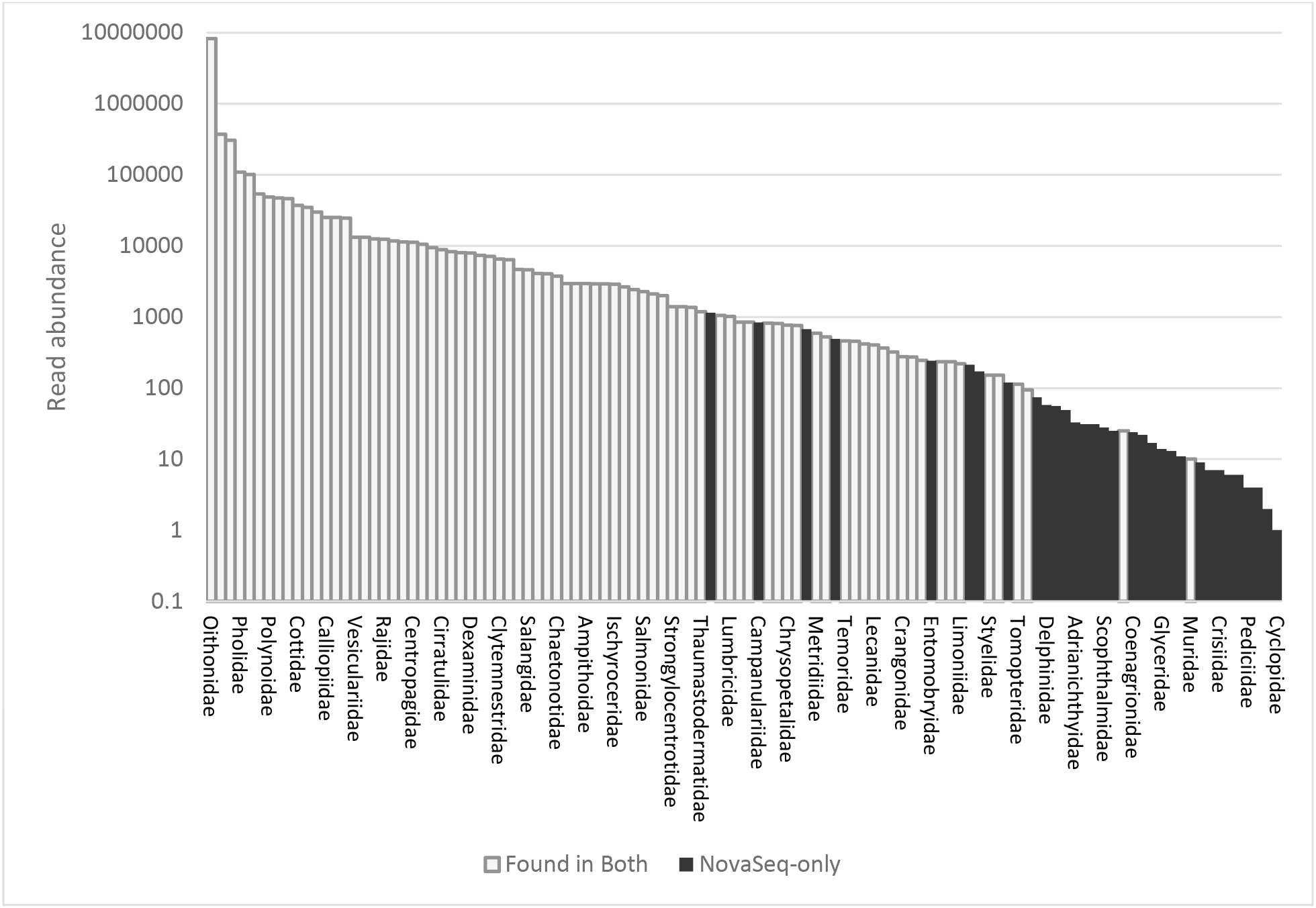
Families detected on the NovaSeq ranked by read count (note the y-axis has a logarithmic scale). The white bars indicate taxa that were detected by both the NovaSeq and the MiSeq, while black bars indicate taxa detected solely by the NovaSeq. We note that most of the taxa missed by the MiSeq have low read abundance.

These findings have great implications for environmental metabarcoding studies and other similar amplicon-based biodiversity analyses: it is possible that the taxa identified through traditional observational approaches but missed through eDNA are a consequence of the instrument used to analyze the eDNA. Moreover, would switching to a more advanced platform (i.e., the NovaSeq) eliminate many of these discrepancies?

### 5.3 Possible causes for the difference between platforms

The NovaSeq has many differences from the MiSeq, including: a 2-colour chemistry rather than 4-colour used by the MiSeq; greatly improved hardware (presumably including better image capture abilities); different signal processing software; and a flow cell that has pre-defined binding spots for target DNA instead of the random lawn used by previous Illumina instruments. We suspect but cannot be certain that this last factor—the flow cell—is a significant cause for the NovaSeq’s superior performance. In MiSeq and most previous Illumina instruments, DNA binds to the flow cell in a random fashion. Therefore, to distinguish one spot on the flow cell where DNA has bound from another, the spots are observed by the instrument for the first 25 rounds of sequencing and at that point clusters are determined ^26^. This works well when performing shotgun sequencing (the primary use of Illumina’s instruments) because the spots can be clearly distinguished from each other thanks to the high level of sequence diversity. However, when performing amplicon-based sequencing the variability from one spot to the next—especially within the first 25 bases which likely covers primer regions—is minimal and this can cause two distinct spots to be merged together. To prevent this from happening, Illumina recommends spiking in PhiX genome ^26^, but unless the proportion of PhiX is very high it’s nearly impossible to prevent similar sequences from sitting near each other on the flow cell. Conversely, the spots that DNA anneal to on the NovaSeq flow cell are pre-defined and known by the instrument’s base calling software, so inferring their location is not necessary and this largely prevents the “overclustering” of low diversity reads.

Notably, the MiSeq runs that we performed for this project used the recommended levels of PhiX and the sequencing run statistics all matched Illumina recommendations. Nevertheless, we still suspect that over-clustering is at least partially responsible for the MiSeq missing out on diversity that the NovaSeq was able to detect.

Our results suggest that the NovaSeq 6000 may be a superior instrument for environmental metabarcoding studies especially in complex biodiversity-rich substrates where heterogenous abundance of taxonomic groups may confound detection. However, while the MiSeq is a relatively inexpensive instrument that could plausibly be obtained by many labs, the NovaSeq is expensive and therefore may be out of reach for many laboratories in the near term. For this reason, we suggest a few approaches that may aid obtaining more comprehensive biodiversity from the MiSeq:

#### Multiplex different markers together on the same flow cell

This is already quite a common practice, albeit frequently for money-saving purposes rather than to prevent over-clustering. In theory, multiplexing several markers (while still maintaining adequate sequencing depth per sample), will lead to greater sequence diversity on the flow cell and will reduce the probability of over-clustering.

#### Use large amounts of PhiX spike-in

PhiX genome fragments also serve to increase the complexity on the flow cell, but many labs try to minimize the amount of PhiX they use because it takes up precious sequencing capacity. Paradoxically though, increasing PhiX may in fact increase the number of quality reads generated. Illumina’s own recommendations range from 5-50% ^27^ although in practice most experiments end up at the lower end of this range.

#### Use phased amplicons

Another approach was suggested by ^28^, who designed overlapping 16s amplicons that they described as “phased amplicon sequencing”. Despite covering the same region of interest, the different sequence composition at the 5’ end of the read reduced over-clustering by the MiSeq—so much so that the number of reads passing quality filters increased by 9-47% in their experiments.

## 6 Conclusions

Biodiversity analysis through genomics has enabled widespread applications from human microbiome studies to environmental assessment and monitoring. With rapid advances in sequencing hardware and computational approaches for data analysis, it is important to determine the impact that sequencing technology and strategy have on the data generated, especially where it may influence biological interpretations and their socio-economic implications. Here, we tackled the issue of sequencing technology and depth on an analysis of biodiversity in seawater through eDNA metabarcoding. Our analysis provides direct evidence of the superior utility of the newly introduced NovaSeq platform for elucidating a more comprehensive biodiversity measurement as compared to the current workhorse, the MiSeq platform. Our results strongly suggest that comprehensive detection of biota from eDNA in a complex environment such as the ocean is possible and will aid supporting scientific/societal endeavours for enhanced biodiversity analysis for people and the planet.

## 7 Acknowledgements

The authors would like to thank Kirk Rees, Captain of the *Abrigo*, for assisting with sample collection. This work was partly funded through a Petroleum R&D Grant from InnovateNL (contract number 5405.2121.101), an award from the Atlantic Canada Opportunities Agency’s Atlantic Innovation Fund (project number 781-37749-207993), and a grant from Petroleum Research Newfoundland and Labrador. Any opinions, findings, and conclusions or recommendations expressed in this publication are those of the authors and do not necessarily reflect the views of Petroleum Research or its members.

## 8 Author contributions

MH conceived the idea for this project and provided scientific oversight in experimental design, data analysis and interpretation; and helped with writing the manuscript; NF designed and executed the sampling protocols; JB aided bioinformatics analyses; AM led the laboratory work; GS performed the data analyses, figure generation, and wrote the manuscript.

## 10 Materials and correspondence

All correspondence and material requests should be addressed to M Hajibabaei

