## Supplementary figures for "Comprehensive biodiversity analysis via ultra-deep patterned flow cell technology: a case study of eDNA metabarcoding seawater"

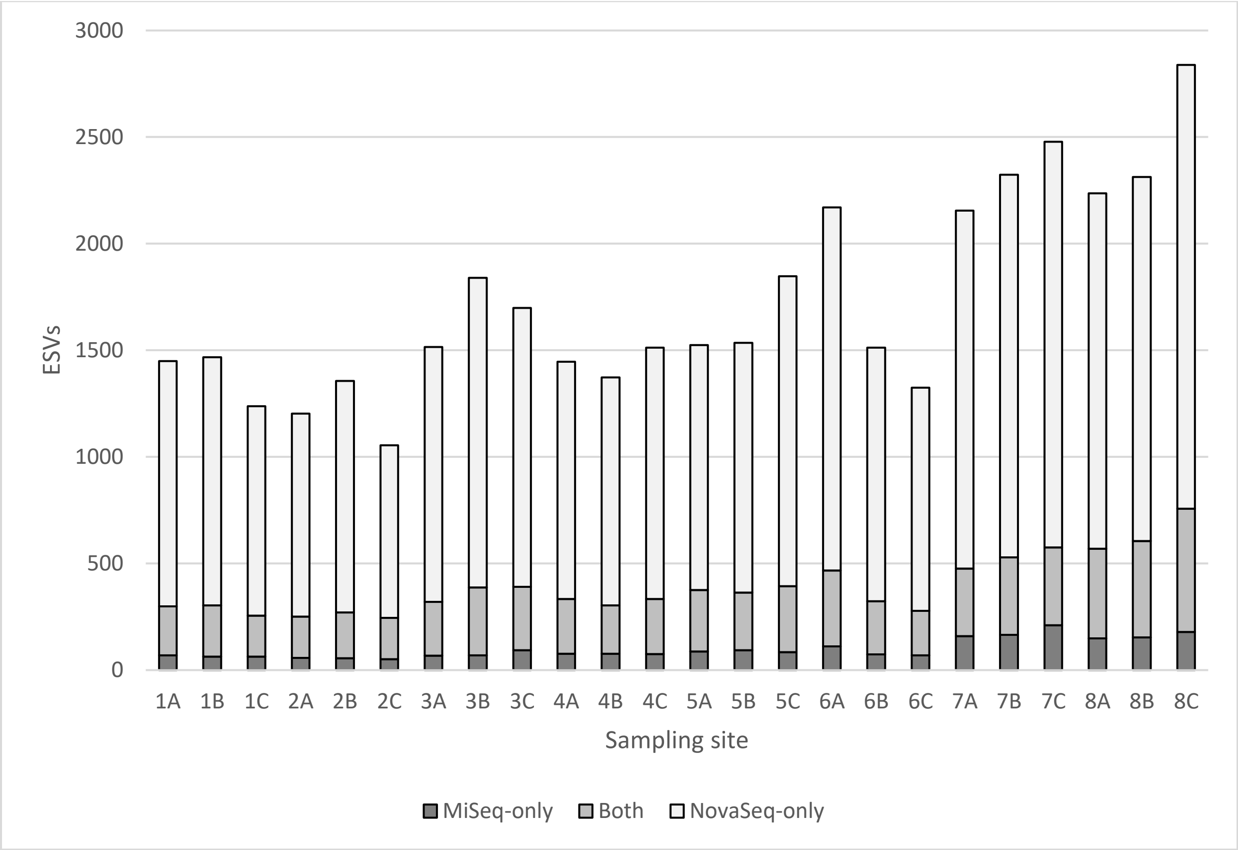


*Figure 1: The NovaSeq finds many more ESVs for the F230 amplicon than the MiSeq, even when correcting for sequencing depth with subsampling. This is true on a replicate-by-replicate and site-by-site basis.*

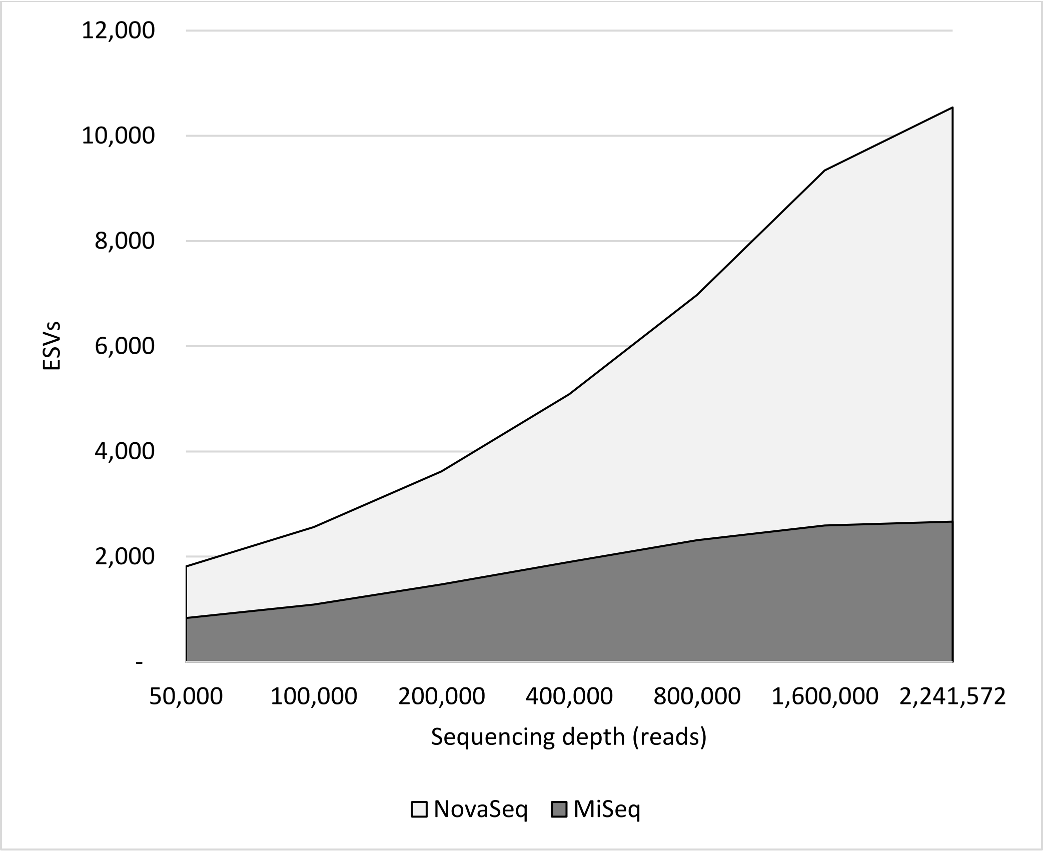


*Figure 2: When all sites/replicates are combined to create an overall accumulation curve for the F230 amplicon, the ability of the NovaSeq to obtain new ESVs is even more pronounced. The number of ESVs detected by the MiSeq seems to level off at approximately 2600 while the NovaSeq has detected more than 10,000 with an upward trajectory indicating more ESVs would be detected with even greater sequencing depth.*

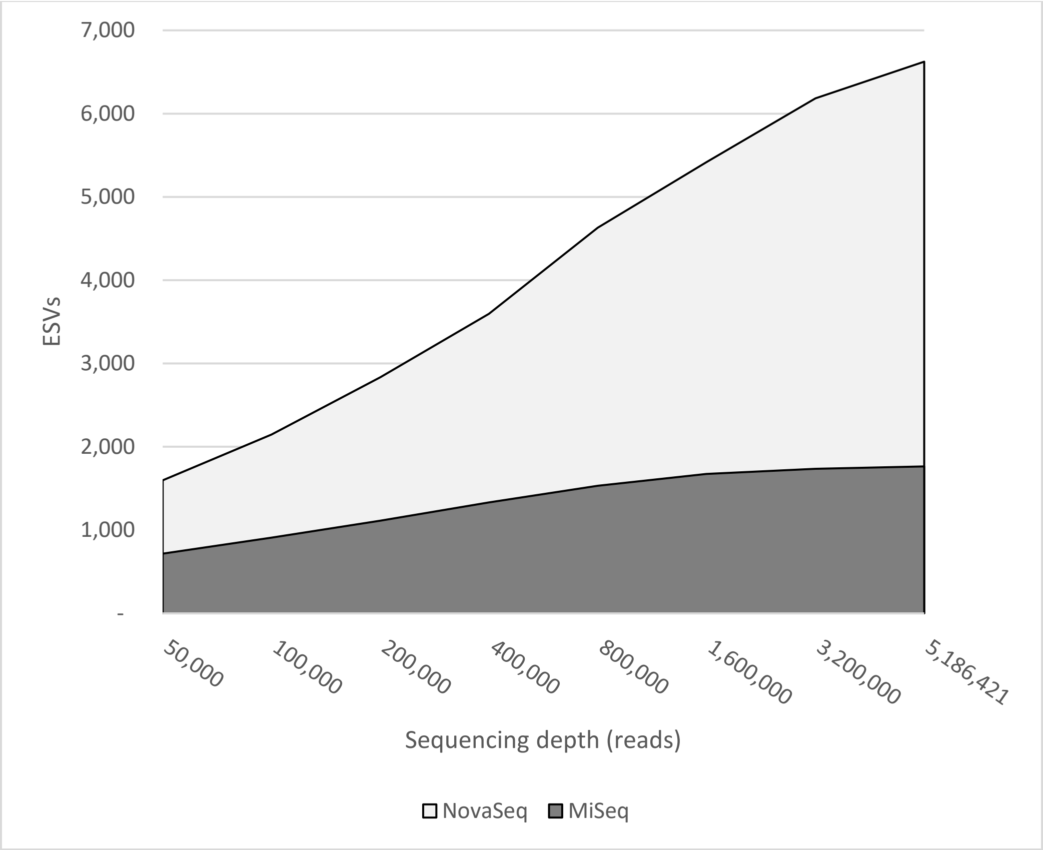


*Figure 3: Even when F230 amplicons are subjected to a very high sequencing depth on the MiSeq, it cannot match the power of the NovaSeq to detect the diversity present in the samples.*
